# External Validation of Machine Learning and EEG for Continuous Pain Intensity Prediction in Healthy Individuals

**DOI:** 10.1101/2023.11.09.566379

**Authors:** Tyler Mari, Jessica Henderson, S. Hasan Ali, Danielle Hewitt, Christopher Brown, Andrej Stancak, Nicholas Fallon

## Abstract

Previous research has predicted subjective pain intensity from electroencephalographic (EEG) data using machine learning (ML) models. However, there is a paucity of externally validated ML models for pain assessment, particularly for continuous pain prediction (e.g., decoding pain ratings on a 101-point scale). We aimed to conduct the first external validation paradigm for ML regression models for pain intensity prediction from EEG data. Ninety-one subjects were recruited across three samples. Sample one (n = 40) was used for model development, sample two (n = 51) was used as a cross-subject external validation set, whilst sample three (n = 25) was used as a within-subjects temporal external validation set. Pneumatic pressure stimuli were delivered to the left-hand index fingernail bed at 10 graded intensity levels. Single-trial time-frequency features of peri-stimulus EEG were used to train a Random Forest (RF) model and long short-term memory (LSTM) network to predict pain intensity responses. Results demonstrated that both the RF model and LSTM network predicted pain intensity significantly more accurately than a random prediction model, with the mean absolute error (MAE) of the RF (best performing model) at 19.59, 21.29, and 18.90 for internal validation, cross-subject external validation, and within-subject external validation, respectively. However, neither model was able to predict pain intensity better than a baseline dummy model, which predicted the mean behavioural rating of the training set and did not have access to neural data. Moreover, in a replication of our recent work, we developed a RF model for the classification of low and high-pain trials, which demonstrated internal and external validation accuracies up to 64% and 58%, respectively. Taken together, our results suggest that using ML and EEG to predict continuous pain ratings is not currently feasible. However, classification models demonstrate some potential, consistently outperforming chance across validation samples. Further improvements such as composite measures are required to elevate ML performance to a clinically meaningful level.

## Introduction

Pain is subjective, complex, and challenging to measure due to an intricate interplay between biological, psychological, and social factors (Bendinger & Plunkett, 2016; Breivik et al., 2008; Gatchel et al., 2007; Younger et al., 2009). The current gold standard of pain assessment is self-report measures, requiring high-level linguistic and social skills, which are unsuitable for individuals who cannot accurately communicate their pain (Herr et al., 2011; Schiavenato & Craig, 2010). Vulnerable populations including non-verbal individuals (Herr et al., 2011; D. Li et al., 2008; McGuire et al., 2016) or individuals with cognitive impairments (Herr et al., 2011; Voepel-Lewis et al., 2002), traumatic brain injury (Arbour & Gélinas, 2014), dementia (Breivik et al., 2008; Herr et al., 2011; Kunz et al., 2009), or disorders of consciousness (Herr et al., 2011; Schnakers & Zasler, 2007), and children (Herr et al., 2011; Witt et al., 2016) are often unable to self-report their pain, which can prevent effective pain management. Therefore, pain assessment techniques that are independent of self-report may facilitate improved pain management in these populations.

Numerous brain regions contribute to pain processing, including the primary (SI) and secondary (SII) somatosensory cortex, insular cortex, anterior and midcingulate cortex, prefrontal cortex, thalamus, amygdala, periaqueductal grey, cerebellum, and brainstem (Duerden & Albanese, 2013; Jensen et al., 2016; Petre et al., 2022; Peyron et al., 2000; Xu et al., 2020). A recent coordinate-based activation-likelihood estimation (ALE) meta-analysis demonstrated consistent pain-related activations independent of stimulus modality, location, and gender in bilateral SII, amygdala, thalamus, brainstem, right middle frontal gyrus, left insula and midcingulate cortex (Xu et al., 2020). The brain regions implicated in pain processing are often considered a distinct pattern, such as the pain matrix or neurologic signature, which exhibits activity changes that encode pain intensity (Garcia-Larrea & Peyron, 2013; Wager et al., 2013). Consequently, neural markers of pain may enable proxy pain assessment.

Electroencephalography (EEG) may demonstrate clinical utility as a proxy pain assessment technique as it is low-cost and easy to use (Mackey et al., 2019; Tivadar & Murray, 2019). Importantly, pain-related changes in cortical oscillations are observable across scalp regions in established frequency bands, which may enable pain assessment (Kim & Davis, 2021; Ploner et al., 2017; Zis et al., 2022). Augmented theta oscillations have been observed during the resting state EEG of individuals with fibromyalgia (Fallon et al., 2018). Additionally, research has demonstrated increased theta amplitudes over central and parietal regions during tactile and painful stimulation, with larger amplitudes observed during painful stimulation (Michail et al., 2016). The contribution of alpha and beta bands in pain processing is well-established, with research consistently demonstrating alpha suppression and beta enhancement during tonic pain stimulation (Chen & Rappelsberger, 1994; Dowman et al., 2008; Huber et al., 2006; Shao et al., 2012). Finally, gamma-band oscillations over SI predict both stimulus and subjective pain intensity (Gross et al., 2007; Zhang et al., 2012). Overall, EEG activity could reliably decode pain intensity.

Supervised machine learning (ML) has been successfully implemented to decode pain-related outcomes using several neuroimaging modalities (Mari et al., 2022; van der Miesen et al., 2019). Specifically, we previously externally validated ML and EEG for low and high pain intensity classification through a multistage validation procedure (Mari et al., 2023). Using 50 time-frequency features consisting of theta, alpha, lower beta, upper beta, and gamma bands from frontal, central, and parietal regions, we classified low and high pain with a cross-validation accuracy of 73.18% using a random forest (RF). Importantly, the model generalised to a novel sample with an accuracy of 68.32%. Further, the model achieved an accuracy of 60.42% on additional external data that used different experimental pain stimulation. Consequently, our results provided robust estimates of ML performance for pain classification on novel samples. However, regression models, which demonstrate finer prediction resolution, should be assessed to improve clinical utility. Obtaining more precise pain estimates may enable improved pain management. For example, predicting continuous ratings enables finer monitoring of pain over time and allows for changes after treatment to be more accurately assessed (e.g., small changes in pain intensity can be identified, which is not possible with broad binary classification; Shirvalkar et al., 2023).

Previous research has used linear regression to predict subjective pain intensity (0-10) from single-trial laser-evoked potentials (LEPs), achieving a mean absolute error (MAE) of 1.03 and 1.82 (lower scores represent better performance) for within- and cross-subject predictions, respectively (G. Huang et al., 2013). Furthermore, Bai and colleagues (Bai et al., 2016) developed a normalisation technique to reduce EEG inter-individual variability and improve model performance, achieving a MAE of 1.17 for cross-subject prediction. Using the same dataset, Li et al. (2018) predicted subjective pain intensity with a MAE of 1.19. Finally, research has demonstrated that subjective pain intensity could be predicted with a MAE of 1.15, using pre- and post-stimulus time-frequency features (Tu et al., 2016). The evidence suggests that EEG and ML can be combined to predict continuous pain ratings.

The previous findings are promising but lack external validation. To comprehensively assess ML performance, models should be evaluated on data that is independent of the training set, as internal validation methods often result in inflated performance metrics (e.g., accuracy) due to overfitting (Cabitza et al., 2021; Lever et al., 2016; Siontis et al., 2015; Vabalas et al., 2019; Varma & Simon, 2006). External validation, which evaluates model performance on novel data obtained from different cohorts, facilities, repositories or collected at a different time, location or using a different experimental paradigm, is essential to obtain robust estimates of model generalisability during prediction model development (Cabitza et al., 2021; Collins et al., 2015). As model performance is often diminished on external data (X. Li et al., 2019; Mari et al., 2023; Siontis et al., 2015), the generalisability and utility of studies that only employ internal validation are unclear and not sufficient evidence to support clinical translation (Bleeker et al., 2003; Ramspek et al., 2021). Although significant further research is required, studies that externally validate pain prediction models are emerging (Furman et al., 2020; Mari et al., 2023).

The present study aimed to externally validate ML and EEG for the prediction of subjective pain intensity both across and within subjects. Firstly, we trained a RF model to predict subjective pain intensity (0-100) using hand-crafted time-frequency EEG features. Secondly, we developed a long short-term memory (LSTM) network to predict subjective pain intensity using the EEG time series from each electrode and frequency band for each trial. Furthermore, model performance was assessed using a multi-stage validation approach consisting of cross-validation (RF model only), internal validation, and external validation (both across and within subjects). The cross-subject validation sample consisted of a new cohort, whilst the within-subject temporal validation sample consisted of participants from the model development sample who completed a second experimental session.

To assess whether the ML models predicted pain intensity better than chance levels, we compared the ML algorithms to two additional dummy models, including a random prediction model and a baseline model that predicted the mean value of all subjective pain responses from the training set. We hypothesised that the RF and LSTM would predict subjective pain intensity (0 – 100) using EEG data more accurately than both the random and baseline models, achieving lower MAE scores for all samples. Secondly, we hypothesised that the LSTM would predict subjective pain intensity more accurately than the RF model on all samples.

## Methods

### Participants

A total of 116 EEG recordings were collected across model development, cross-subject validation, and within-subject temporal validation samples. Participants were recruited using an opportunity sampling method. The model development sample consisted of 40 participants (22 female; 7 left-handed) aged between 18 and 52 (Mean = 27.70 years, standard deviation [SD] = 7.43). The cross-subject validation sample consisted of an additional 51 participants (34 female; 6 left-handed) aged between 19 and 60 (Mean = 27.63 years, SD = 9.65). There was no participant overlap between the development and cross-subject validation samples. Moreover, a total of 25 participants aged between 21 and 53 (14 female; 4 left-handed; Mean = 28.96 years, SD = 8.01) from the development sample completed the study for a second time after a minimum of 12 weeks had elapsed from their first session (Mean = 108.68 days, SD = 10.92), resulting in a within-subject temporal validation sample. Participants were at least 18 years old, had normal or corrected-to-normal vision, no neurological conditions, no acute pain at the time of participating, no history of chronic pain and no injuries to the left-hand index finger that may affect sensory perception (e.g., nerve damage). Participants provided written informed consent before participation and all methods were conducted in compliance with the Declaration of Helsinki. This research received ethical approval from the University of Liverpool Health and Life Sciences Research Ethics Committee. Participation was reimbursed at a rate of approximately £13.33 per hour. The raw data is available from authors on reasonable request.

### Pneumatic Pressure Stimulator

Tonic pain stimulation was delivered to the fingernail bed of the left-hand index finger using a custom-built pneumatic pressure stimulator (Dancer Design, St. Helens, UK), as utilised in previous pain research from our lab and others (Mari et al., 2023; Watkinson et al., 2013). The pneumatic stimulator consisted of a pneumatic force controller, which directed air from an 11.1-L aluminium cylinder into the stimulator. This lowered a 1cm^2^ probe to deliver the desired force. The stimulator was controlled using a LabJack U3 printed circuit board for interface. The pressure was mechanically limited to a maximum of 3.5 bar (12kg/cm^2^) to reduce the risk of injury.

### Procedure

The experiment was conducted in the EEG laboratory at the University of Liverpool between June and October 2022. On arrival at the lab, participants were seated 1 metre away from a 23-inch 1080p LCD monitor inside a Faraday cage. Participants received a verbal description of the experiment before reading the information sheet and providing written consent. A custom mould of the participant’s left-hand index finger, which correctly positioned and maintained the finger underneath the stimulator probe, was created using a two-part silicone elastomer. The stimulator probe was aligned to stimulate the fingernail bed of the left-hand index finger. Additionally, participants were offered foam earplugs (28dB) to minimise any potential noise. Following alignment, participants underwent a thresholding procedure to identify their maximum-intensity stimulus.

Before the initiation of the thresholding block, participants were instructed to rate the pain intensity of each stimulus on a 101-point (0-100) numerical rating scale by using the mouse in their right hand to select the desired point on the scale. Scale anchoring was set at 0 which represented no sensation, and 100 which reflected extreme pain. Additionally, 30 represented the pain threshold and was denoted on the rating scale with the number 30 and the term “pain threshold”. The scale rating included vertical bars which denoted increments of 10. Participants were informed that ratings below 30 represented non-painful stimulation (e.g., touch) and that a rating of 0 indicated that they did not feel the probe, or the probe did not touch their finger due to finger compression. Participants were also informed that the stimulus intensity that elicited a rating of 70 or above would be used as their maximum intensity and that this value reflected upper moderate pain intensity.

A staircase procedure was implemented for the thresholding section. Here, the pressure intensity was initialised at 0.4 bar and increased in steps of 0.2 bar whilst the participant’s rating was below 40. Once the rating exceeded 40, the increment was reduced to 0.1 bar. The maximum intensity of the stimulus was limited to 3.0 bar (10.5kg/cm^2^) using custom software. The participants initialised the block by pressing the space bar, which began lowering the probe. During this period, a black fixation cross was presented on the screen. For each trial, the pressure stimulus had a rise time of 1 second, which reflected the time taken for the probe to go from 0 bar to the desired intensity. Subsequently, the trial intensity was maintained for 3 seconds, released, and followed by a 4-second wait period. The rating scale was then displayed until the participant successfully rated their pain intensity. Following the rating phase, the scale was replaced with a black fixation cross, and a further two-second wait time was implemented. The thresholding block was terminated once the participant had rated one of the stimulus intensities as at least 70 representing upper moderate pain. The pressure that elicited this rating was set as the maximum intensity. Moreover, participants were informed that the intensity could be adjusted throughout the session if it was either too painful or not painful enough. Stimuli intensity changes were set at fixed values of 0.1, 0.2, or 0.3 bar depending on the judgment of the researcher (e.g., an inspection of the pain ratings) and discussions with the participant. Following the completion of the thresholding procedure, participants exited the Faraday Cage for the EEG cap fitting.

For the main experimental block, a set of 10 stimuli intensities was created for each participant. The 10 stimuli intensities were linearly spaced between the minimum pressure (0.4 bar) and the upper-pressure limit selected for the participant (e.g., the pressure that elicited a pain intensity rating ≥ 70 on the thresholding procedure block). In each block, a total of 40 pseudo-randomised stimuli were delivered, which consisted of four randomised repetitions of each of the 10 stimuli intensities (see Stimuli Randomisation Procedure). Before the start of the block, the participant’s finger was realigned under the stimulator probe and occluded from sight. Subsequently, participants were provided with verbal and written instructions for the task.

Each trial consisted of a baseline period, stimulus delivery phase, and post-stimulus rating segment. A baseline period of 4 seconds was implemented, followed by a 1-second stimulus rise time, where the stimulus intensity increased from 0 bar to the desired stimulus intensity in 1/10^th^ increments every 0.1 seconds. Once the stimulus had reached the desired intensity, it was maintained for a total of 3 seconds before being released. Following the end of the stimulation period, a 4-second post-stimulus phase was implemented. A black fixation cross was presented on the centre of the screen continuously during the previous segments.

Following the completion of the stimulation period in each trial, participants were required to rate their subjective pain intensity using the same 101-point scale as the thresholding procedure. However, participants were informed that they could report any subjective pain intensity that corresponded to their experience on that trial, i.e., that rating above the previous 70 threshold was permitted without the experiment terminating. The rating scale section had an infinite duration with a minimum of 2 seconds. Following the rating phase, the fixation cross was presented on the screen and a subsequent forced wait period was conducted. Each trial had a minimum inter-trial interval of 16 seconds. Following the completion of the experimental task, the EEG cap was removed and participants completed the pain catastrophising scale (Sullivan et al., 1995) before being debriefed and reimbursed for their time.

The experiment consisted of four blocks, resulting in a total of 160 stimuli, with each block lasting approximately 15 minutes. Each block was separated by a distractor task, resulting in a minimum inter-block interval of 5 minutes. All experimental procedures were delivered using PsychoPy (Peirce, 2007).

### Stimuli Randomisation Procedure

Stimuli were pseudo-randomised using a custom randomisation algorithm. Pseudo-randomisation was conducted to prevent the clustering of high-intensity stimuli, which could have become too painful for the participant. Consequently, the 10 stimulus intensities were ordered from minimum to maximum and arranged into 5 pools that contained two adjacent stimuli. For example, pool one included the two lowest stimuli intensities, whilst pool 5 contained the two highest stimuli intensities. An empty array was created to store the final stimuli ordering. Subsequently, one of the 5 pools was chosen at random and one value in that pool was selected, removed, and added to the final ordering array. If the selected pool was the highest intensity pool, on the next iteration, the algorithm was forced to select from pools one, two, or three which contained stimuli at approximately low and moderate intensities. If the maximum intensity pool was not selected, the algorithm could select any of the pools that still contained values on the next iteration. This process was repeated until the 10 stimuli were shuffled and appended to the final ordering list. This process was repeated 4 times for each block, resulting in a total of 40 stimuli. To prevent high stimuli clustering at the end and beginning of each iteration, additional safety measures were implemented. On the second, third, and fourth iterations, the algorithm was prevented from selecting one of the highest-intensity pools first if the last intensity from the previous iteration was also high, which further prevented high-intensity stimuli from grouping. This process was conducted for each block and participant, meaning that all blocks across all participants had a unique ordering.

### EEG Acquisition

EEG recordings were continuously obtained using a 129-channel EGI System (Electrical Geodesic Inc., EGI, now Magstim EGI, Eugene, Oregon, USA) and a sponge-based Geodesic sensor net. The correct net position was achieved by aligning the net with respect to three anatomical landmarks: two pre-auricular points and the nasion. Electrode-to-skin impedances were monitored and maintained below 50 kΩ for all electrodes throughout the experiment. A recording bandpass filter was set at 0.001 – 200 Hz, whilst the sampling rate was set at 1000 Hz. Finally, Cz was used as the reference electrode.

### EEG Pre-processing

Automatic EEG pre-processing, using the Harvard Automated Processing Pipeline for Electroencephalography (HAPPE; version 3; Gabard-Durnam et al., 2018), was conducted due to the large sample size of this study. Line noise at 50 (± 2) Hz was removed using CleanLine (Mullen, 2012), high- and low-pass filters of 0.5 and 70 Hz were applied, and the data were resampled to 500 Hz. Subsequently, bad channels that did not contain useable brain data (e.g., channels affected by excessive movement) were then interpolated and the remaining data underwent wavelet thresholding artefact correction. The data were then segmented into trial epochs with a period of -4 seconds to 6 seconds relative to the stimulus onset. Any channels with remaining artefacts were interpolated and segment rejection was employed for unusable segments. Here, automated epoch rejection was conducted using two criteria: the amplitude range, which was set at -150 to 150 mV as recommended by HAPPE authors (Gabard-Durnam et al., 2018) and segment similarity. Trial epochs identified in this process were marked for rejection. Following the trial rejection, the data were re-referenced using an average reference (Lehmann, 1987). Finally, the data was visually inspected. From the visual inspection procedure, it was identified that the data still contained excessive line noise. Consequently, an additional notch filter (50 ± 2 Hz) was applied to the data to remove any remaining line noise. Table 1 shows the average remaining number of trials for each of the 10 stimuli intensities after pre-processing for all three samples. Following pre-processing, a total of 81.12%, 80.70%, and 81.53% of trials were retained in the model development, cross-subject validation, and within-subject validation samples, respectively. There were no significant differences between the remaining number of trials for each stimuli intensity in the model development sample (p = .153), cross-subject validation sample (p = .818), or within-subject temporal validation sample (p = .876).

**Table 1.** Average number and percentage of trials retained per subject after EEG pre-processing.

| Stimulus Intensity | Sample |  |  |
| --- | --- | --- | --- |
|  | Model Development | Cross-subject | Within-subject |
| 1 | 12.78 ± 1.99 (80%) | 12.71 ± 1.85 (79%) | 13.20 ± 1.73 (83%) |
| 2 | 12.98 ± 2.04 (81%) | 12.92 ± 1.84 (81%) | 12.92 ± 1.75 (81%) |
| 3 | 13.05 ± 1.85 (82%) | 13.14 ± 1.93 (82%) | 13.24 ± 1.67 (83%) |
| 4 | 12.90 ± 2.25 (81%) | 12.84 ± 1.67 (80%) | 13.08 ± 1.44 (82%) |
| 5 | 13.08 ± 1.99 (82%) | 12.86 ± 1.88 (80%) | 13.20 ± 2.08 (83%) |
| 6 | 13.28 ± 1.84 (83%) | 12.63 ± 2.16 (79%) | 13.28 ± 1.74 (83%) |
| 7 | 13.18 ± 1.99 (82%) | 13.02 ± 1.84 (81%) | 13.04 ± 1.79 (82%) |
| 8 | 13.53 ± 1.99 (85%) | 12.96 ± 1.77 (81%) | 12.76 ± 2.09 (80%) |
| 9 | 12.60 ± 1.86 (79%) | 12.96 ± 1.78 (81%) | 12.68 ± 1.89 (79%) |
| 10 | 12.60 ± 1.98 (79%) | 13.08 ± 1.72 (82%) | 13.04 ± 1.77 (82%) |

Spectral analysis was conducted using MATLAB 2020b (The MathWorks, Inc., Natick, Massachusetts, USA) and EEGLAB 2021.1 (Delorme & Makeig, 2004). The power spectral density (PSD) was estimated using Welch’s method. The PSD was calculated for each trial from -4 seconds to 6 seconds relative to the onset stimulus onset, in 1-second segments, shifted in 0.01-second increments. The data were smoothed using 7 multi-taper Slepian sequences. The PSD was calculated between 1 and 70 Hz, with a resolution of 1 Hz. Relative band power changes were calculated across each time point and frequency in the trial epoch using the event-related desynchronisation (ERD) method (Pfurtscheller & Aranibar, 1979; See Equation below). The ERD estimate at each datapoint (A in the equation) is computed by subtracting the mean PSD of the baseline period (-3.5 to -0.5; R) followed by a subsequent numerical transformation to express the relative change in power as a percentage value.

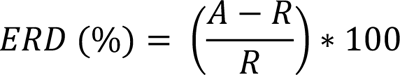

Negative ERD values reflect band power decreases in the active period, relative to the baseline segment, which indicates cortical activation (Neuper & Pfurtscheller, 2001; Pfurtscheller & Aranibar, 1977; Pfurtscheller & Lopes da Silva, 1999; Pfurtscheller & Neuper, 1992). Positive ERD values represent band power increases, which generally reflect cortical inhibition and are referred to as event-related synchronisation (ERS; Pfurtscheller, 1992, 2001). The data were transformed into established frequency bands which are distinct in spatiotemporal dynamics and functional associations: theta (4 – 7 Hz), alpha (8 – 12 Hz), lower beta (16 – 24 Hz), upper beta (25 – 32 Hz) and gamma (33 – 70 Hz) for the ML analysis (Keil et al., 2022; Schomer & Lopes, 2010). This was achieved by averaging the time series within the boundaries of the five frequency bands. Topographical maps, to illustrate power changes from baseline to low and high experimental pain stimulation conditions are reported in the results section. ERD visualisation was conducted and reported following recommendations from previous research (Pfurtscheller & Aranibar, 1977, 1979) and is consistent with our previous work (Mari et al., 2023).

### Machine Learning Procedure

The model development, cross-subject, and within-subject validation samples were processed using the same pipeline but were handled separately to prevent data leakage which could have compromised the external validation procedure (Luo et al., 2016). Firstly, we conducted feature engineering by computing candidate predictors from the single-trial time-frequency transformed data. Eighteen candidate features were calculated for ERD outputs across the five frequency bands and 128 electrodes, resulting in a total of 11,520 potential features. The features were primarily comprised of descriptive statistics of the relative band power changes in each frequency band. The features were calculated from the active segment of the trial window [0-5.5s]. Fifteen of the features were identical to our previous research (see Mari et al., 2023), with the number of peaks, number of troughs, and peak-to-trough ratio also included in this study in an attempt to provide the model with additional features to improve classification performance. All features were calculated using in-built MATLAB functions, where possible, and are consistent with previous research (Anuragi & Sisodia, 2020; Mari et al., 2023; Sai et al., 2019; Vargas-Lopez et al., 2021; Vimala et al., 2019).

Single-trial EEG is significantly hampered by noise and inter-trial variability (Faisal et al., 2008; Kaplan et al., 2005; Marathe et al., 2014). Consequently, an outlier interpolation procedure was employed as outliers do not follow clear patterns, which impairs ML performance (Maniruzzaman et al., 2018). Additionally, interpolation was implemented to maximise the dataset size, as smaller datasets are at an increased risk of overfitting (Vabalas et al., 2019). Outliers were identified and replaced for each participant through linear interpolation using the *filloutliers* MATLAB function, in line with our previous research (Mari et al., 2023). Outliers were identified as values that exceeded three median absolute deviations. Interpolated values were calculated from neighbouring non-outlier data points. A total of 6.46 ± 0.67%, 6.39 ± 0.76%, and 6.33 ± 0.38% of the data were interpolated for the development sample, cross-subject validation sample, and within-subjects temporal validation sample, respectively.

The data were subsequently processed for ML using Python and Scikit-learn (Pedregosa et al., 2011), with the random seed value set as 123. Firstly, the model development sample was split into a cross-validation and an internal holdout test set. Here, 10% (4 participants) of the model development sample was selected for the holdout validation set. The participants within the holdout validation set were randomly selected from a subset of participants (n = 15) from the model development sample, who did not participate in the within-subject temporal validation sample. The features for each validation set were scaled to between 0 and 1 and univariate feature selection was implemented. Once the features had been successfully ranked, a custom sequential feature selection procedure was conducted. Firstly, a baseline RF regressor was developed using only the highest-ranking feature, with no hyperparameter optimisation. This was conducted as RF models demonstrate good performance with default parameters and require minimal hyperparameter tuning (Bentéjac et al., 2021; Fernández-Delgado et al., 2014). Here, we used stratified k-fold validation (k=10) for the cross-validation procedure. Subsequent features were sequentially added to the model until a maximum of 70 features had been included. Seventy was selected as the limit for the number of features as this represented 1/70^th^ of the total of cross-validation sample observations (4685 trials). The limit was implemented to reduce model complexity (e.g., the curse of dimensionality), whilst providing enough features to enable successful ML training. The model and features that achieved the best cross-validation performance were selected as the final feature set for the full model development procedure. Through univariate feature selection, a total of 55 features demonstrated optimal cross-validation performance.

After identifying the optimal features, the final RF regressor was developed and trained on the development sample dataset. A RF model was selected as research has shown that RFs provide optimal real-world performance and are robust to overfitting (Dong et al., 2020; Fernández-Delgado et al., 2014; Jiang et al., 2020; Mienye & Sun, 2022). Additionally, the RF model achieved the best performance in our previous work (Mari et al., 2023). Random search with a maximum of 50,000 iterations was conducted for hyperparameter optimisation, which evaluated a range of lower and upper bounds for hyperparameter values to identify the optimal configuration (Bergstra & Bengio, 2012; Géron, 2019; Yang & Shami, 2020). Stratified k-fold validation was integrated into the hyperparameter optimisation. A value of k = 10 was used for this study. Hyperparameter optimisation was only conducted in relation to cross-validation performance, the hold-out, cross-subject, and within-subject validation sets did not inform model development. Following hyperparameter optimisation, the model was refit to the entire training set. The final model was subsequently evaluated on the internal hold-out validation and external validation sets.

### Time Series ML

In addition to the RF model, we also developed a bidirectional LSTM network for time-series prediction. LSTMs are a type of recurrent neural network that can identify and learn long-term dependencies in sequence input data through the use of memory cells, which can remember inputs across time steps (Hochreiter & Schmidhuber, 1997; LeCun et al., 2015). The regulation of the stored information (e.g., when to remove it) is managed by gating mechanisms, such as forget gates (Gers et al., 2000; Hochreiter & Schmidhuber, 1997; LeCun et al., 2015). Bidirectional LSTMs provide an extension of the original LSTM architecture, by extracting information in both forwards and backwards directions. Essentially, two LSTMs are developed and trained using the input data, with one of the networks being trained on the original time series and the second being trained backwards on the time series. This allows the network to learn relationships in either direction.

We developed a 7-layer LSTM network which had a total of 2.4 million learnable parameters. The input layer dimensions were 640×550, with 640 input time-series, which reflected the 5 frequency bands (theta, alpha, lower beta, upper beta, and gamma) for each of the 128 scalp electrodes, by 550 time points. Subsequently, a bidirectional LSTM layer with 256 hidden units was implemented. On all LSTM layers, the state activation was the hyperbolic tangent function, whilst a sigmoid was used as the gate activation function. Furthermore, a dropout layer with a probability of 0.2 was implemented to help reduce the risk of overfitting, which was followed by an additional bidirectional LSTM layer, with 128 hidden units. A final dropout layer, again with a 0.2 probability, was implemented prior to the fully connected layer. Finally, the data was passed through a regression layer to obtain a continuous prediction. The network was trained for a maximum of 200 epochs, with a learn rate of 0.001, and a minibatch size of 64 samples. Gradient clipping was implemented with a threshold of 1. Finally, the network that achieved the best validation loss was selected as the final model.

### ML Evaluation

The primary performance metric in this study was the mean absolute error (MAE) which is consistent with previous research (Bai et al., 2016; G. Huang et al., 2013; L. Li et al., 2018; Tu et al., 2016). The MAE represents the average error between the true label and the predicted label (Mari et al., 2022; Willmott & Matsuura, 2005) and can be calculated using the following equation:

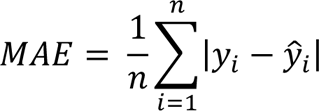

Where y_i_ is the true label and ŷ_i_ is the model predicted value.

For each validation set (internal hold-out, cross-subject, and within-subject validation), we calculate the MAE for each subject and average across the validation set to obtain both individual and sample accuracies.

Unlike binary classification, there is no obvious chance level to ascertain whether the models outperform random prediction. Therefore, we developed two additional dummy models that allowed us to evaluate the effectiveness of the ML models. Dummy models provide a way of establishing a minimum expected baseline performance by using simple heuristics to provide predictions without any knowledge of the input features (Fontana et al., 2019; Johnson et al., 2016; Moon et al., 2020). For example, dummy models in classification may always predict the majority class, whilst they may predict the mean of the outcome variable in the training set for regression tasks (Fontana et al., 2019; Johnson et al., 2016; Moon et al., 2020). Simple heuristics can often outperform ML performance, so the use of dummy models provides an effective baseline to evaluate the effectiveness of ML models. This process ensures that the models are not exploiting simple rules instead of learning from the input features (Fontana et al., 2019).

Firstly, we developed a dummy model that predicted a random number between 0 and 100 for each trial and calculated the MAE for each validation set. This was termed the random model. Secondly, we developed a baseline (dummy) model that predicted the mean value of subjective response data from all trials in the training set. The baseline model provided a measure of the accuracy of predictions that can be made using only behavioural pain ratings (e.g., omitting neural data). To evaluate which models achieved the lowest MAE, we conducted three one-way ANOVAs that assessed each of the four models (random, baseline, RF, LSTM) using IBM SPSS 27 (IBM Corp., Armonk, New York, USA). Bonferroni correct post-hoc tests were conducted to investigate significant main effects.

## Results

### Subjective Pain Ratings

The average pain ratings for each stimulus intensity across all three samples is reported in Table 2. Three simple linear regressions were conducted using SPSS 27 (IBM Corp., Armonk, New York, USA) to assess the relationship between stimulus intensity and subjective pain intensity in the development sample, cross-subject validation, and within-subject temporal validation sample, respectively. For the development sample, the regression model was significant and predicted 77% variance (R^2^ = .77, F (1,5196) = 17531.63, p < .001). Stimulus intensity was a significant positive predictor of subjective pain intensity (b = 7.40, se = 0.06, p <.001, 95% CI = 7.29 to 7.51). For the cross-subject validation sample, the regression model was also significant and predicted 75% variance (R^2^ = .75, F (1,6583) = 19538.91, p < .001). Stimulus intensity was a significant positive predictor of subjective pain intensity (b = 7.38, se = 0.05, p <.001, 95% CI = 7.28 to 7.49). Finally, the regression model was significant and predicted 73% variance (R^2^ = .73, F (1,3259) = 8986.29, p < .001) for the within-subjects validation sample. Again, stimulus intensity was a significant positive predictor of subjective pain intensity (b = 6.60, se = 0.07, p <.001, 95% CI = 6.46 to 6.74). Figure 1 illustrates the relationship between stimulus intensity and subjective pain intensity for all three samples.

**Table 2.** Descriptive statistics for each stimulus intensity and sample.

| Stimulus Intensity | Model Development |  |  | Cross-subject<br>Validation |  |  | Within-subject<br>Validation |  |  |
| --- | --- | --- | --- | --- | --- | --- | --- | --- | --- |
|  | Mean | SD | N | Mean | SD | N | Mean | SD | N |
| 1 | 4.52 | 5.62 | 511 | 4.62 | 7.62 | 648 | 8.35 | 8.60 | 330 |
| 2 | 9.17 | 8.46 | 519 | 9.10 | 9.40 | 659 | 13.46 | 10.05 | 323 |
| 3 | 13.87 | 9.50 | 522 | 14.84 | 10.62 | 670 | 17.92 | 10.62 | 331 |
| 4 | 21.82 | 12.42 | 516 | 22.93 | 12.50 | 655 | 24.55 | 12.14 | 327 |
| 5 | 29.93 | 13.15 | 523 | 30.58 | 13.54 | 656 | 31.33 | 12.25 | 330 |
| 6 | 38.05 | 13.22 | 531 | 39.32 | 14.38 | 644 | 39.17 | 12.97 | 332 |
| 7 | 46.14 | 14.03 | 527 | 47.18 | 14.70 | 664 | 45.36 | 12.82 | 326 |
| 8 | 55.06 | 12.62 | 541 | 55.11 | 14.09 | 661 | 53.50 | 11.82 | 319 |
| 9 | 61.11 | 11.56 | 504 | 61.37 | 12.75 | 661 | 59.55 | 11.56 | 317 |
| 10 | 67.82 | 10.53 | 504 | 67.82 | 10.97 | 667 | 65.97 | 9.99 | 326 |

**Figure 1.**
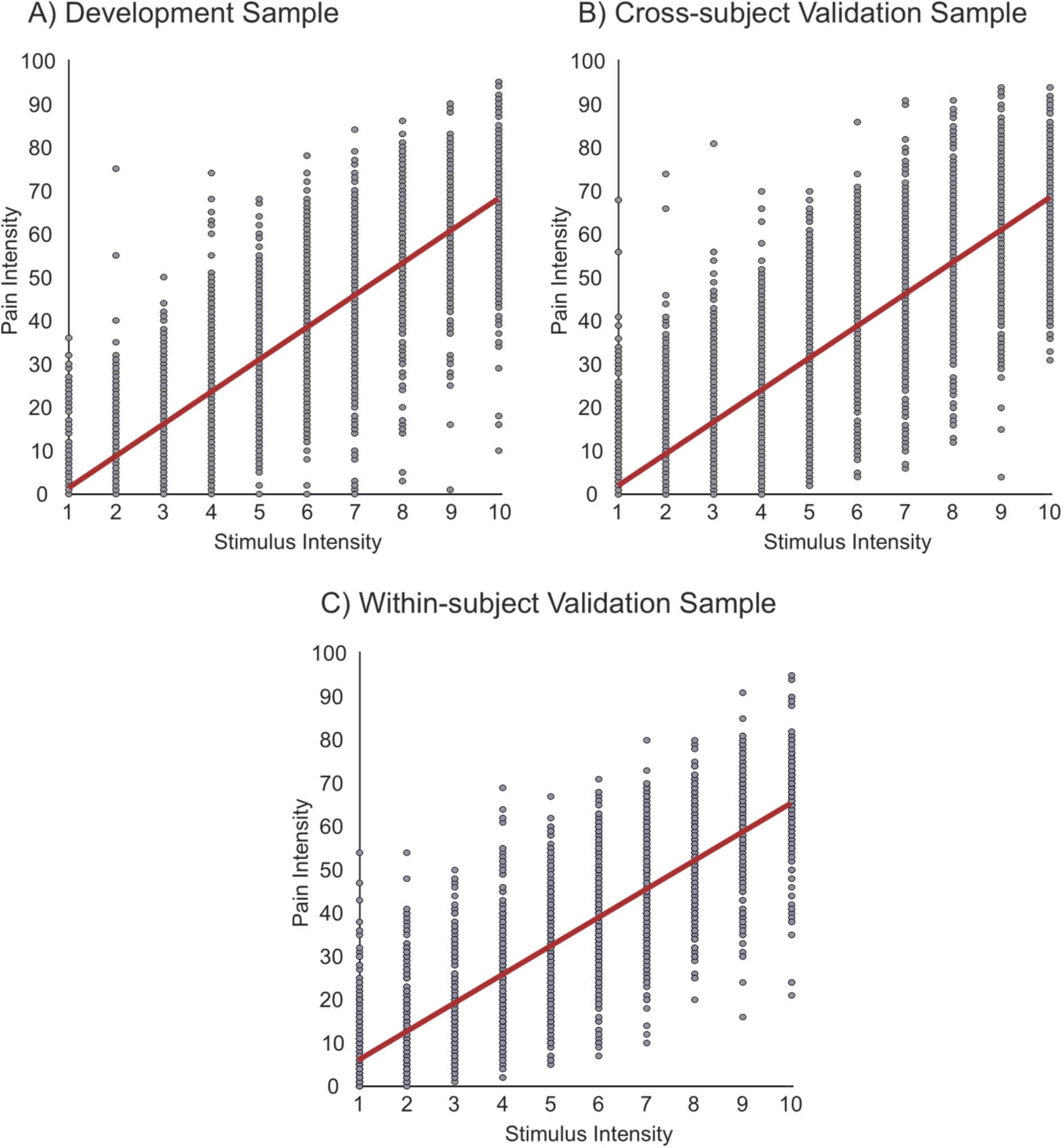
The relationship between stimulus intensity and subjective pain intensity for the model development sample (A), Cross-subject validation sample (B), and Within-subject validation sample (C). The red line represents the least squares regression line.

### ERD/S

Topographic maps illustrating the difference between low and high pain conditions in the 91 unique participants are shown in Figure 2. The figure shows the time-frequency changes during the rest (-3 – -2 s relative to the onset of stimulation) and the active period (1 – 2 s relative to the onset of stimulation). The active period represents the first second of maximum pressure following the completion of the stimulation rise time. Topographic plots demonstrating relative band power changes in frequency bands Theta (4 – 7Hz), Alpha (8 – 12Hz), Lower Beta (16 – 24Hz), Upper Beta (25 –32Hz), and Gamma (33 – 70Hz) are reported. The left pair of columns represent the rest and active periods for the low pain condition, whilst the right pair of columns represent the high pain condition.

**Figure 2.**
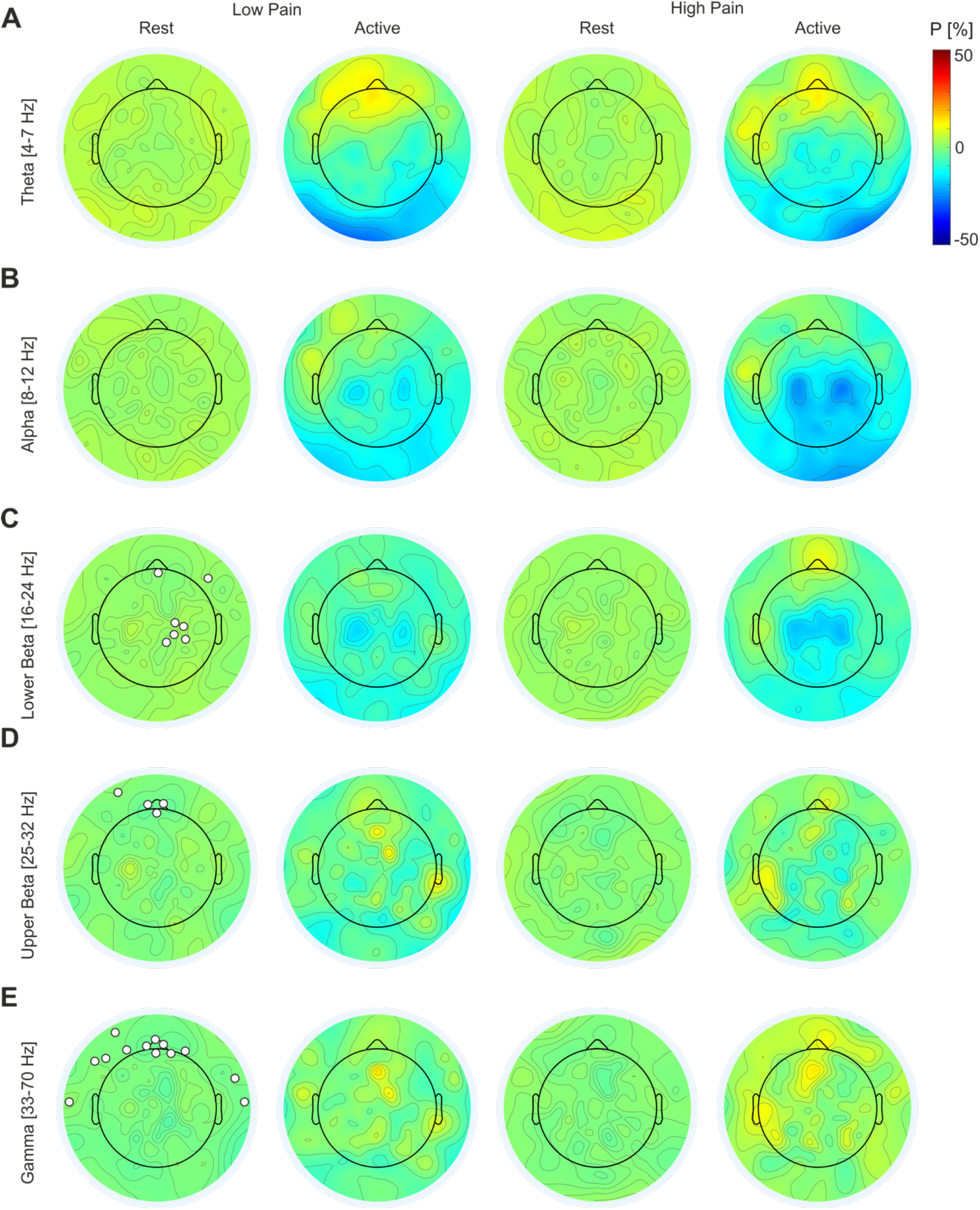
Grand average band power changes during rest (-3 – -2s) and active pressure stimulation (1 – 2 s) from all 91 unique participants (combined sample one and two). The trial period spanned from -4 s to 6 s relative to the trial onset, with a baseline period of -3.5 s to - 0.5 s. The active period was selected in line with previous recommendations (Pfurtscheller & Aranibar, 1977, 1979) and represented 1 s of continued pressure immediately after the stimulator reached the desired stimulus intensity level. The topographic maps show the band power changes in low and high pain intensity conditions and from rest to active periods in Theta (A), Alpha (B), Lower Beta (C), Upper Beta (D), and Gamma (E). The white circles represent the electrode locations of the features used in the ML classification of low and high pain trials. P = percentage power change from baseline.

In the Theta band, we observed ERS over frontal electrodes in both low and high pain conditions (Figure 2A). Moreover, sensorimotor, and occipital ERD was also observed during both low and high pain conditions in theta band. There was strong bilateral ERD in the alpha band, observed over sensorimotor regions in both low and high pain conditions (Figure 2B). Here, the intensity of bilateral alpha ERD was clearly enhanced during the high pain condition, relative to low pain. Bilateral ERD was also observed in both the lower and upper beta bands over sensorimotor regions (Figure 2C/D), but the pattern was visibly weaker in the upper Beta band when compared to the lower Beta band. In both lower and upper Beta bands, a similar pattern to alpha processing was observed, with stronger ERD evident in the high pain condition. Finally, for the Gamma band power changes, we observed ERS over frontal regions in both low and high pain stimulation, which appeared to be more widespread during the high pain stimulation condition (Figure 2E).

### Machine Learning Results

The final number of observations for each validation set/sample is presented in Table 3. From the feature selection procedure, a total of 55 features were optimal. The features were distributed across scalp regions and frequency bands. The theta and alpha features were predominantly located over a frontal-central electrode. Features in the beta band were located predominantly over frontal regions, with additional features calculated from peripheral electrodes. The gamma band provided most features (approximately 50%) and was distributed over frontal regions and central-parietal regions, respectively.

**Table 3.** The number of observations for each validation set.

| Validation Set | Number of Trials |
| --- | --- |
| Training Sample (n = 36) | 4685 |
| Internal Hold-out Validation (n = 4) | 513 |
| Cross-subject (n = 51) | 6585 |
| Within-subject (n = 25) | 3261 |
| Total | 15044 |

The MAE for all the models and validation sets are reported in Table 4. For the validation sample, the RF model predicted pain intensity on a 101-point scale with a MAE of 19.59 points, whilst the LSTM demonstrated similar performance with an average error of 19.97.

**Table 4.** The MAE for each model across all validation sets.

|  | Internal Validation |  |  |  | External Validation |  |  |  |
| --- | --- | --- | --- | --- | --- | --- | --- | --- |
|  | Cross-Validation |  | Hold-out Validation |  | Cross-subject |  | Within-subject |  |
|  | Mean | SD | Mean | SD | Mean | SD | Mean | SD |
| RF | 20.49 | 0.69 | 19.59 | 2.30 | 21.29 | 3.40 | 18.90 | 3.34 |
| LSTM | - | - | 19.97 | 2.50 | 21.17 | 3.53 | 19.03 | 2.73 |
| Random Model | - | - | 32.61 | 3.24 | 33.41 | 2.90 | 32.34 | 2.76 |
| Baseline Model | - | - | 19.98 | 2.58 | 21.19 | 3.51 | 19.08 | 3.19 |

Optimal RF hyperparameters: Number of estimators = 145, Maximum depth = 8, Minimum samples to split = 19, Minimum samples at leaf = 12, Maximum features = sqrt, Bootstrap = True.

Whereas the random model and baseline model predicted subjective pain intensity with a MAE of 32.61 and 19.98, respectively. A one-way ANOVA demonstrated that there was a significant main effect of model on the MAE (F (3,12) = 22.74, p < .001, ƞ_p_^2^ = .85). Bonferroni post-hoc tests showed that the random model had a significantly higher error for predicting subjective pain intensity than the RF model (p < .001), LSTM network (p < .001), and the baseline model (p < .001). However, there were no significant differences between the baseline model and the RF model (p = 1.00) or the LSTM network (p = 1.00) for pain intensity prediction. Finally, there was no significant difference between the RF model and the LSTM network (p = 1.00).

Regarding the cross-subject validation sample, the LSTM demonstrated the most accurate pain intensity prediction, achieving a MAE of 21.17. Here, the RF predicted subjective pain intensity with an error of 21.29, which was less accurate than the baseline model, which demonstrated a MAE of 21.19. Finally, the random model demonstrated an error of 33.41. A one-way ANOVA demonstrated a significant main effect of ML model on the MAE for the cross-subject validation sample (F (3,200) = 169.12, p < .001, ƞ_p_^2^ = .72). In line with the internal validation results, the random prediction model had significantly greater error for pain predictions when compared to the RF model (p < .001), LSTM network (p < .001), and the baseline model (p < .001). Again, there were no significant differences between the baseline model and the RF (p = 1.00) or the LSTM network (p = 1.00). No significant difference was observed between the RF and the LSTM (p = 1.00).

Finally, the RF demonstrated the most accurate predictions for pain intensity for the within-subject temporal validation sample, achieving a MAE of 18.90. The LSTM was the next best-performing model, achieving an error of 19.03. Finally, the random model and baseline model demonstrated less accurate predictions, achieving MAEs of 32.24 and 19.08, respectively. The ANOVA demonstrated a significant main effect of model on the MAE (F (3,96) = 121.86, p < .001, ƞ_p_^2^ = .79). The random model demonstrated significantly larger prediction error than the RF model (p < .001), LSTM network (p < .001), and the baseline model (p < .001). No further significant differences were observed between the baseline model and RF model (p = 1.00), the baseline model and LSTM network (p = 1.00), or the RF model and LSTM network (p = 1.00).

Calibration for regression is not as clearly calculated as classification (Levi et al., 2022). Moreover, calibration assessment is only required when the models demonstrate good predictive capability (Alba et al., 2017). Therefore, as neither the LSTM nor the RF demonstrated predictive performance that surpassed the baseline model, we did not formally assess calibration. However, to provide insight into the behaviour of the model, we aimed to visualise the model’s predicted values and the true pain intensity values. We only provide the visualisations for the RF as, on average, it demonstrated the most accurate predictions. Figure 3 illustrates the relationship between subjective pain intensity and the RF model’s predicted values for all three validation sets. Briefly, the results show that the RF model tended to predict values in the middle of the scale, with most of the predictions for all samples falling between 25 and 50. Moreover, there is not a clear trend between the predicted values and true values, suggesting that the model failed to extract relevant information to provide informative predictions. Finally, the model failed to predict extreme values (e.g., < 25 and > 65), meaning that the model could not accurately predict the lowest and highest intensities.

**Figure 3.**
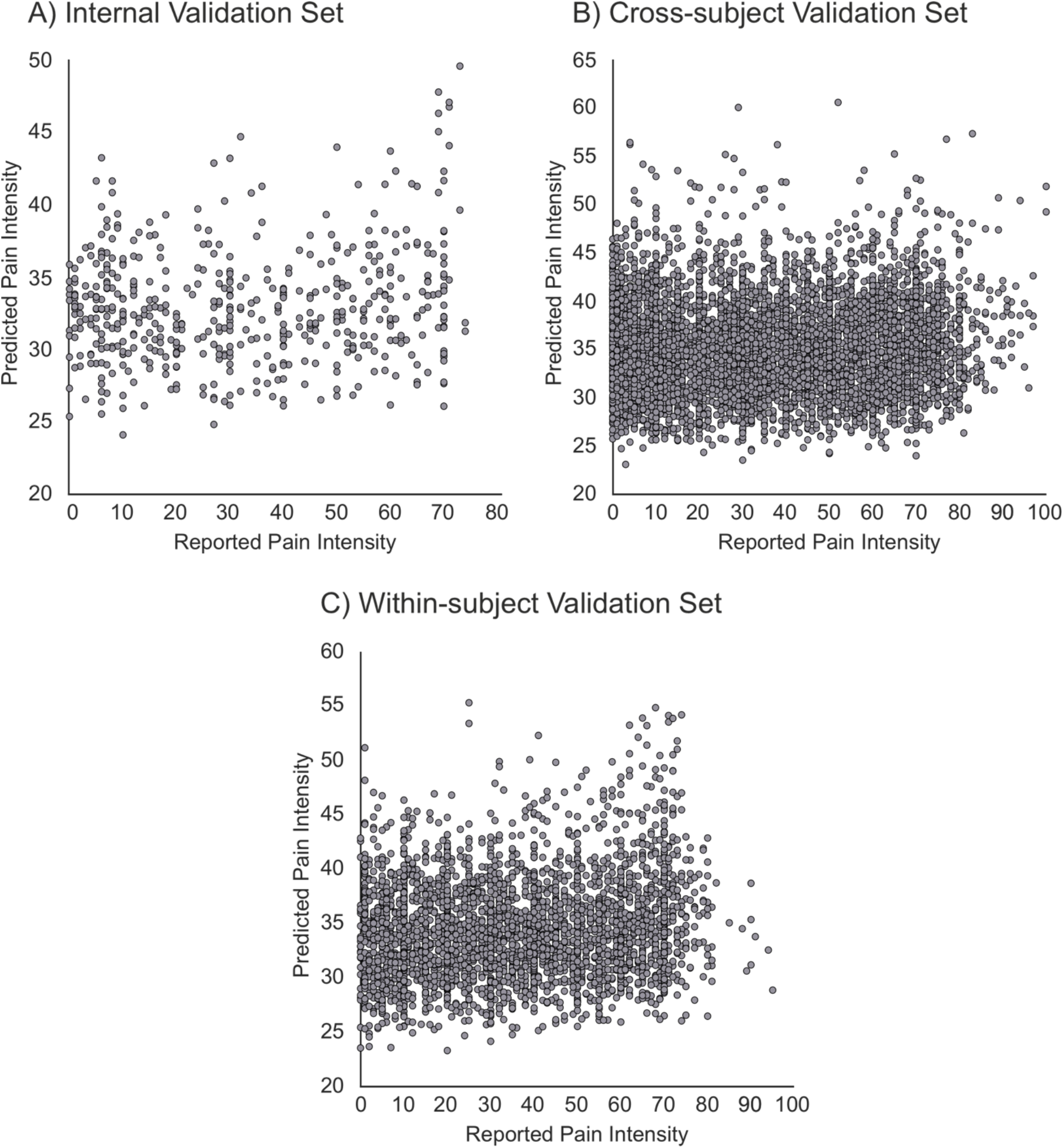
The relationship between the RF model’s predicted pain intensity and the reported subjective pain intensity for the internal validation set (A), cross-subject validation set (B), and within-subject validation set (C).

### Exploratory Analysis

As the regression model was unable to outperform the baseline model and provide informative predictions, we attempted to develop a binary classification RF model for the prediction of low and high pain trials, providing a theoretical replication of our prior work (Mari et al., 2023). This was achieved by selecting stimulus intensities that were comparable to the low and high stimuli intensities of our previous study. Stimuli intensities 4 and 5 were chosen for the low intensity, whilst 9 and 10 were chosen for the high intensity set, as these elicited comparable subjective pain ratings to our earlier model development sample (Mari et al., 2023). The RF model was developed and evaluated using a similar procedure to the regression model for feature selection and hyperparameter optimisation. Several common performance metrics were used to evaluate the performance of the model. Here, we report the accuracy, AUC, Brier score, F1 score, precision, and recall, which is consistent with existing research (Mari et al., 2022, 2023). The ML classification results are reported in Table 5. Moreover, the optimal feature locations for the classification are displayed in Figure 2.

**Table 5.** Performance metrics for binary RF model for the prediction of low and high pain intensity.

|  | Accuracy | SD | AUC | SD | Brier | SD | F1 | SD | Precision | SD | Recall | SD |
| --- | --- | --- | --- | --- | --- | --- | --- | --- | --- | --- | --- | --- |
| **Internal Validation** |  |  |  |  |  |  |  |  |  |  |  |  |
| Cross-validation | 0.64 | 0.05 | 0.67 | 0.06 | 0.23 | 0.01 | 0.61 | 0.08 | 0.66 | 0.05 | 0.58 | 0.11 |
| Hold-out | 0.62 | 0.11 | 0.68 | 0.18 | 0.22 | 0.05 | 0.58 | 0.12 | 0.61 | 0.15 | 0.57 | 0.15 |
| **External Validation** |  |  |  |  |  |  |  |  |  |  |  |  |
| Cross-subject | 0.57 | 0.08 | 0.62 | 0.10 | 0.24 | 0.03 | 0.52 | 0.13 | 0.60 | 0.11 | 0.50 | 0.20 |
| Within-subject | 0.58 | 0.09 | 0.62 | 0.11 | 0.25 | 0.03 | 0.49 | 0.19 | 0.59 | 0.15 | 0.47 | 0.24 |

Optimal RF hyperparameters: Number of estimators = 2280, Maximum depth = 15, Minimum samples to split = 3, Minimum samples at leaf = 7, Maximum features = log2, Bootstrap = True.

The results can be separated into internal and external validation procedures. For internal validation, the classification results demonstrated that the RF model outperformed the theoretical chance level (50%) for the classification of low and high-pain intensity trials using EEG features, achieving a cross-validation accuracy and AUC of 64% and 0.67, respectively. Similar results were observed for the holdout validation sample, with the RF model exceeding chance level performance. The results demonstrated that the RF classified subjective pain intensity with an accuracy of 62% and an AUC of 0.68.

Regarding external validation, the results show that the RF model exceeded chance performance on both the cross-subject and within-subject validation samples. Here, for the cross-subject validation sample, subjective pain intensity was classified with an accuracy of 57% and an AUC of 0.62. Similar results were observed for the within-subject validation sample, with the model producing an accuracy of 58% and an AUC of 0.62. Overall, the RF demonstrated similar, above-chance performance, for both external validation datasets.

Finally, we also assessed calibration as the model performance exceeded chance performance. Calibration refers to the agreement between the model’s predicted outcome value and the true outcome value (Alba et al., 2017; Luo et al., 2016; Van Calster et al., 2019). Calibration curves, which illustrate the relationship between the predicted probabilities (x-axis) and the observed probabilities (y-axis), are the preferred way to assess binary classification models (Moons et al., 2015; Van Calster et al., 2016, 2019). To achieve this, we split the data into 10 equal bins, which represented the probabilities between 0 and 1 (Y. Huang et al., 2020). The calibration curves for the holdout validation, cross-subject external validation, and within-subject external validation samples are presented in Figure 4. To interpret the calibration curves, when the model performance line is above the reference line (which represents perfect calibration), it suggests that the model underestimates the probability of the incidence. Whereas a model line below the reference line indicates that the model is overestimating the probability of the incidence. The calibration results suggest that the predictions from the RF model are reasonably well calibrated. There are instances of the model underestimating the incidence of the event at the lower probabilities and overestimating the higher probabilities. However, the calibration curves typically follow the expected trend, suggesting that the model provides reasonably accurate probability estimates.

**Figure 4.**
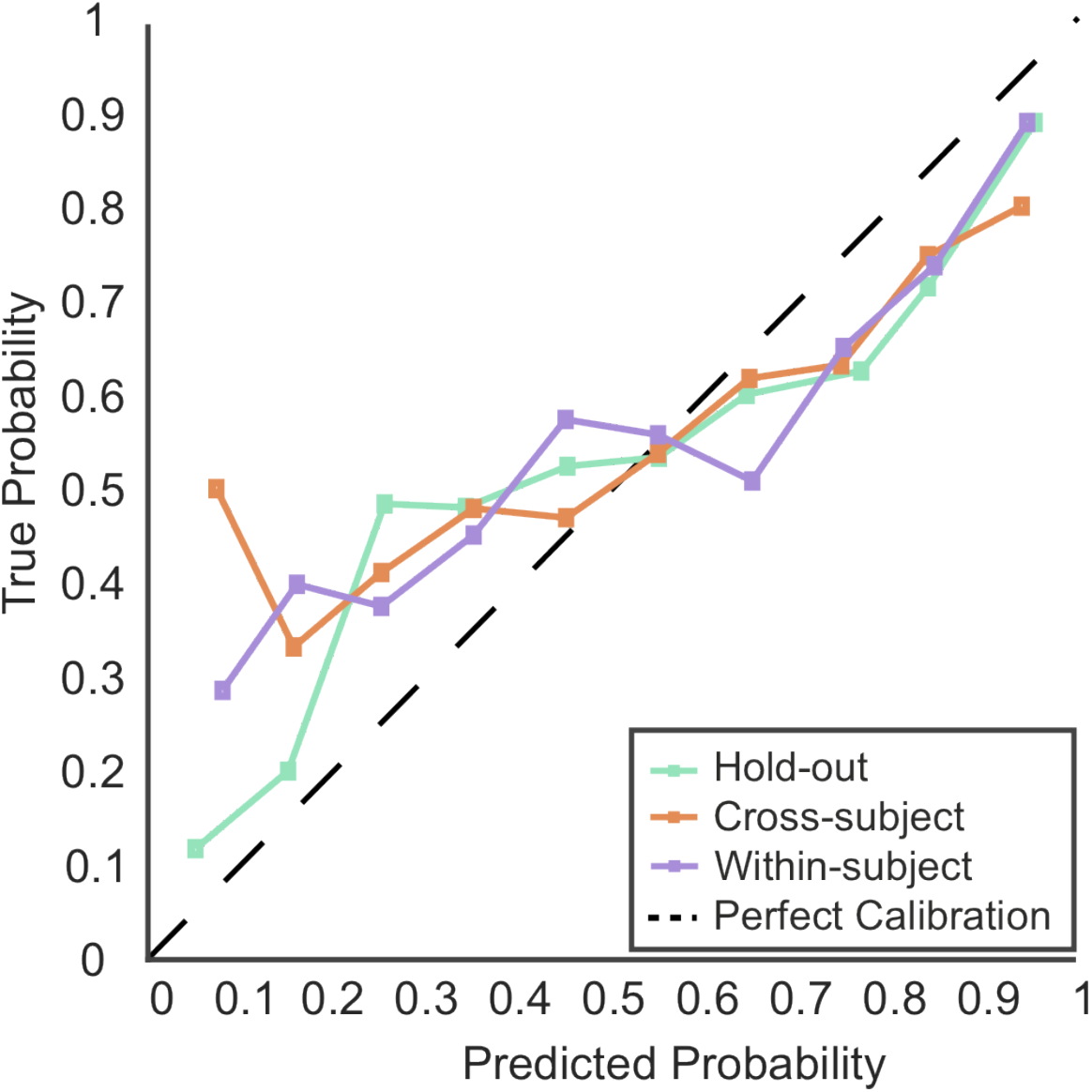
Calibration curves for the internal holdout validation sample, and two external validation samples. The black dotted line 45° represents perfect calibration.

## Discussion

We aimed to externally validate ML for the prediction of subjective pain intensity both across and within subjects using single-trial EEG. We hypothesised that both the RF model and LSTM network would predict pain intensity more accurately than a random prediction model and a baseline prediction model across three validation sets. The results partially support the hypotheses, with both the RF and LSTM demonstrating more accurate pain intensity predictions than the random model on all validation datasets. However, the baseline (dummy) model which utilised only behavioural data also outperformed the random model. Moreover, neither the RF nor the LSTM outperformed the baseline model on any of the validation sets. Our second hypothesis that the LSTM would outperform the RF was not supported as no differences in model performance were observed on any of the validation sets. The results suggest that regression models trained on oscillatory EEG data cannot predict subjective pain intensity more accurately than simple heuristics using the current approach.

As the regression models failed to provide meaningful predictions, we subsequently aimed to replicate our previous research by classifying low and high-pain intensity trials (Mari et al., 2023). Here, the RF model accurately classified the conditions with better-than-chance accuracies of 64%, 62%, 57%, and 58% for cross-validation, internal holdout validation, cross-subject external validation, and within-subject external validation, respectively. Moreover, the model demonstrated AUCs of 0.67, 0.68, 0.62, and 0.62 for the validation samples. Overall, the classification results are promising but require improvement to demonstrate clinically meaningful levels (e.g., AUC ≥ 0.75 Fan et al., 2006).

Despite not exceeding baseline performance, our regression metrics are comparable to previous research. Huang and colleagues (2013) demonstrated that continuous pain ratings could be predicted using LEPs with a MAE of 1.03 and 1.82 (11-point scale) for within-subject and cross-subject predictions, respectively. Further studies report MAEs of approximately 1.2 for continuous pain prediction using ML and EEG (Bai et al., 2016; L. Li et al., 2018; Tu et al., 2016). Therefore, our findings are comparable to existing literature. However, the previous results are not externally validated and should be interpreted cautiously (Cabitza et al., 2021; Siontis et al., 2015; Vabalas et al., 2019; Varma & Simon, 2006). Consequently, the results of this study provide the most robust estimates of the potential of ML and EEG for pain intensity prediction, providing realistic estimates of both model performance and clinical potential.

Our classification results support existing literature that has demonstrated that EEG and ML can classify low and high pain trials with cross-validated accuracies between 62 and 89.58% (Bai et al., 2016; G. Huang et al., 2013; Mari et al., 2023; Okolo & Omurtag, 2018; Schulz et al., 2012; Tu et al., 2016). In addition, the external validation results are comparable to our previous work which demonstrated that ML could predict low and high pain trials across subjects using two external datasets, previously achieving accuracies of 68% and 60%, respectively (Mari et al., 2023). Despite observing slightly reduced performance in the current study, the results support the potential of EEG and ML for pain intensity classification.

This study highlights the challenges associated with predicting continuous pain intensity using EEG and ML, whilst supporting the potential of classification models. Due to the large sample size and multi-stage external validation, our results provide robust performance estimates of the effectiveness of ML and EEG for pain intensity prediction. However, the increased rigour may explain the observed performance reduction compared to our previous research (Mari et al., 2023). Small samples demonstrate increased performance variability (Arbabshirani et al., 2017; Vabalas et al., 2019; Varoquaux, 2018). Varoquaux (2018) reviewed ML performance across several domains including Alzheimer’s, autism, and depression, and identified that model performance decreased as a function of sample size. Furthermore, small external validation samples can also result in imprecise model performance estimates (Snell et al., 2021). As much of the previous research consists of small samples, performance is likely inflated due to increased variability (Mari et al., 2022). The impact of small samples may also explain the observed minor reduction in external validation performance compared to our recent research (Mari et al., 2023), which comprised fewer participants. Therefore, this study supports our previous work with improved robustness to give increased confidence in the findings. Overall, ML and EEG remain promising for pain classification, but improved performance from robustly designed studies remains imperative. Finally, whilst continuous pain intensity prediction is desirable for finer prediction resolution which would enable improved pain assessment and treatment monitoring/recommendations (Shirvalkar et al., 2023), it appears unrealistic within the current approach.

The current study has several limitations. Firstly, EEG has a low signal-to-noise ratio (Tivadar & Murray, 2019), which likely affects the ability of the ML algorithm to extract meaningful patterns at the single trial level. Single-trial EEG is inherently noisy due to the variability and volatility of neural activity and due to the physical limitations of the apparatus (Faisal et al., 2008; Kaplan et al., 2005; Marathe et al., 2014; Tivadar & Murray, 2019). The diminished signal quality may have impaired model performance. In future, methods to improve the signal-to-noise ratio should be explored (e.g., spatial filtering; Miao et al., 2021; Rivet et al., 2009). Spatial filtering aims to increase the signal-to-noise ratio by maximising the differences between two classes and increasing discriminability (Blankertz et al., 2008; Miao et al., 2021; Rivet et al., 2009). However, spatial filtering techniques are also prone to overfitting (Blankertz et al., 2008; Grosse-Wentrup et al., 2009), which could exacerbate an already prevalent issue in this research area. Therefore, spatial filtering techniques should be applied cautiously to minimise overfitting risk.

Moreover, whilst the RF classification model demonstrated reasonably well-calibrated predictions, there were instances where the model provided inaccurate estimates. The calibration assessment demonstrated that the model estimates were occasionally too extreme, as the models occasionally tended to underestimate the lower probabilities and overestimate the higher probabilities, which indicates a degree of overfitting (Van Calster et al., 2019). However, as single-trial EEG is an inherently noisy signal (Faisal et al., 2008; Kaplan et al., 2005; Marathe et al., 2014; Tivadar & Murray, 2019), it is expected that the model captures some random noise in the training set. Therefore, we recommend interpreting the results with caution. However, the calibration of the present study was arguably improved compared to our previous research (Mari et al., 2023), which demonstrates the improved robustness of this study. Moreover, as calibration is rarely assessed (Christodoulou et al., 2019; Mari et al., 2022), the calibration assessment of this study is an area of novelty and represents a methodical improvement over the existing literature. Nevertheless, future research should improve model calibration through techniques such as Platt scaling or isotonic regression (Y. Huang et al., 2020).

Despite promising classification results, further developments are imperative for clinical translation. Research has suggested that binary classification models with an of AUC less than 0.75 are not clinically meaningful (Fan et al., 2006). Therefore, future research should prioritise improving model performance on external data towards clinically meaningful results. The use of composite biological measures as predictive of subjective pain intensity may improve performance (Rockholt et al., 2023; Tracey et al., 2019). Methods such as heart rate variability, skin conductance, and pupillometry have shown promise for pain assessment (Cowen et al., 2015). For example, skin conductance has been shown to discriminate the presence or absence of pain in postoperative patients with promising sensitivity and specificity (Ledowski et al., 2006). Moreover, recent research has shown that ML, electrodermal activity, photoplethysmography, and respiration measures could predict the presence or absence of pain with accuracies up to approximately 94% (Fernandez Rojas et al., 2023), although this finding was not externally validated. In future, the potential clinical utility of a combination of several physiological measurements for pain classification should be explored.

Alternative neuroimaging techniques may also prove advantageous for improving model performance. Magnetencephalography (MEG) may be a potential candidate. MEG records magnetic fields as opposed to electrical fields of EEG, which are not distorted by anatomical features (e.g., differences in skull thickness), resulting in improved signal-to-noise ratio and spatial localisation of signals (Singh, 2014). Therefore, MEG may improve signal quality and consequently model performance. However, MEG is more expensive and less practical than EEG (Singh, 2014) which reduces the potential clinical applications, as EEG is easily applied and low-cost (Mackey et al., 2019; Tivadar & Murray, 2019). Although MEG is being used clinically in the surgical treatment of epilepsy (Singh, 2014), currently, there is limited research using MEG and ML for pain classification. One recent study demonstrated that ML and MEG could classify healthy controls and chronic migraine patients with an accuracy greater than 86% (AUC > 0.9). Moreover, the study also demonstrated that ML could correctly classify patients into either chronic migraine and episodic migraine, as well as chronic migraine or fibromyalgia patients with high accuracies (Hsiao et al., 2022). Moreover, given the ongoing development of low-cost MEG that can operate at room temperature (Boto et al., 2017), future research should investigate the utility of MEG for pain intensity prediction, perhaps combining it with EEG for multimodal imaging (Singh, 2014; Yoshinaga et al., 2002), which may further improve performance.

## Conclusion

Our results suggest that prediction of fine-graded resolution pain intensity, such as categorising individual subjective experience on a graded scale, may not be possible using the current approach, as model performance including EEG data did not exceed a simple baseline model. Despite this, our results remain promising for predicting discrete categories of pain (e.g., low, and high pain), with the RF model exceeding chance performance for both cross-subject and within-subject predictions using external data. Due to the large sample size, the current study provides the most robust estimates for the potential of ML and EEG for pain intensity classification. Increasing the signal-to-noise ratio of EEG remains a priority, whilst composite measures should be explored and externally validated in future research to improve the performance of ML algorithms. Overall, ML and EEG can accurately predict discrete levels of subjective pain intensity, but performance levels are not currently sufficient to be considered clinically meaningful or to support the development of translation tools.

## References

Alba, A. C., Agoritsas, T., Walsh, M., Hanna, S., Iorio, A., Devereaux, P. J., McGinn, T., & Guyatt, G. (2017). Discrimination and Calibration of Clinical Prediction Models. JAMA, 318(14), 1377. 10.1001/jama.2017.12126

Anuragi, A., & Sisodia, D. S. (2020). Empirical wavelet transform based automated alcoholism detecting using EEG signal features. Biomedical Signal Processing and Control, 57, 101777. 10.1016/j.bspc.2019.101777

Arbabshirani, M. R., Plis, S., Sui, J., & Calhoun, V. D. (2017). Single subject prediction of brain disorders in neuroimaging: Promises and pitfalls. NeuroImage, 145, 137–165. 10.1016/j.neuroimage.2016.02.079

Arbour, C., & Gélinas, C. (2014). Behavioral and Physiologic Indicators of Pain in Nonverbal Patients with a Traumatic Brain Injury: An Integrative Review. Pain Management Nursing, 15(2), 506–518. 10.1016/j.pmn.2012.03.004

Bai, Y., Huang, G., Tu, Y., Tan, A., Hung, Y. S., & Zhang, Z. (2016). Normalization of pain-evoked neural responses using spontaneous EEG improves the performance of EEG-based cross-individual pain prediction. Frontiers in Computational Neuroscience, 10(APR). 10.3389/fncom.2016.00031

Bendinger, T., & Plunkett, N. (2016). Measurement in pain medicine. BJA Education, 16(9), 310–315. 10.1093/bjaed/mkw014

Bentéjac, C., Csörgő, A., & Martínez-Muñoz, G. (2021). A comparative analysis of gradient boosting algorithms. Artificial Intelligence Review, 54(3), 1937–1967. 10.1007/s10462-020-09896-5

Bergstra, J., & Bengio, Y. (2012). Random search for hyper-parameter optimization. Journal of Machine Learning Research, 13(2), 281–305.

Blankertz, B., Tomioka, R., Lemm, S., Kawanabe, M., & Muller, K. (2008). Optimizing Spatial filters for Robust EEG Single-Trial Analysis. IEEE Signal Processing Magazine, 25(1), 41–56. 10.1109/MSP.2008.4408441

Bleeker, S. ., Moll, H. ., Steyerberg, E. ., Donders, A. R. ., Derksen-Lubsen, G., Grobbee, D. ., & Moons, K. G. . (2003). External validation is necessary in prediction research: Journal of Clinical Epidemiology, 56(9), 826–832. 10.1016/S0895-4356(03)00207-5

Boto, E., Meyer, S. S., Shah, V., Alem, O., Knappe, S., Kruger, P., Fromhold, T. M., Lim, M., Glover, P. M., Morris, P. G., Bowtell, R., Barnes, G. R., & Brookes, M. J. (2017). A new generation of magnetoencephalography: Room temperature measurements using optically-pumped magnetometers. NeuroImage, 149, 404–414. 10.1016/j.neuroimage.2017.01.034

Breivik, H., Borchgrevink, P. C., Allen, S. M., Rosseland, L. A., Romundstad, L., Breivik Hals, E. K., Kvarstein, G., & Stubhaug, A. (2008). Assessment of pain. British Journal of Anaesthesia, 101(1), 17–24. 10.1093/bja/aen103

Cabitza, F., Campagner, A., Soares, F., García de Guadiana-Romualdo, L., Challa, F., Sulejmani, A., Seghezzi, M., & Carobene, A. (2021). The importance of being external. methodological insights for the external validation of machine learning models in medicine. Computer Methods and Programs in Biomedicine, 208, 106288. 10.1016/j.cmpb.2021.106288

Chen, A. C. N., & Rappelsberger, P. (1994). Brain and Human pain: Topographic EEG amplitude and coherence mapping. Brain Topography, 7(2), 129–140. 10.1007/BF01186771

Christodoulou, E., Ma, J., Collins, G. S., Steyerberg, E. W., Verbakel, J. Y., & Van Calster, B. (2019). A systematic review shows no performance benefit of machine learning over logistic regression for clinical prediction models. Journal of Clinical Epidemiology, 110, 12–22. 10.1016/j.jclinepi.2019.02.004

Collins, G. S., Reitsma, J. B., Altman, D. G., & Moons, K. (2015). Transparent reporting of a multivariable prediction model for individual prognosis or diagnosis (TRIPOD): the TRIPOD Statement. BMC Medicine, 13(1), 1. 10.1186/s12916-014-0241-z

Cowen, R., Stasiowska, M. K., Laycock, H., & Bantel, C. (2015). Assessing pain objectively: the use of physiological markers. Anaesthesia, 70(7), 828–847. 10.1111/anae.13018

Delorme, A., & Makeig, S. (2004). EEGLAB: an open source toolbox for analysis of single-trial EEG dynamics including independent component analysis. Journal of Neuroscience Methods, 134(1), 9–21. 10.1016/j.jneumeth.2003.10.009

Dong, X., Yu, Z., Cao, W., Shi, Y., & Ma, Q. (2020). A survey on ensemble learning. Frontiers of Computer Science, 14(2), 241–258. 10.1007/s11704-019-8208-z

Dowman, R., Rissacher, D., & Schuckers, S. (2008). EEG indices of tonic pain-related activity in the somatosensory cortices. Clinical Neurophysiology, 119(5), 1201–1212. 10.1016/j.clinph.2008.01.019

Duerden, E. G., & Albanese, M.-C. (2013). Localization of pain-related brain activation: A meta-analysis of neuroimaging data. Human Brain Mapping, 34(1), 109–149. 10.1002/hbm.21416

Faisal, A. A., Selen, L. P. J., & Wolpert, D. M. (2008). Noise in the nervous system. Nature Reviews Neuroscience, 9(4), 292–303. 10.1038/nrn2258

Fallon, N., Chiu, Y., Nurmikko, T., & Stancak, A. (2018). Altered theta oscillations in resting EEG of fibromyalgia syndrome patients. European Journal of Pain, 22(1), 49–57. 10.1002/ejp.1076

Fan, J., Upadhye, S., & Worster, A. (2006). Understanding receiver operating characteristic (ROC) curves. CJEM, 8(01), 19–20. 10.1017/S1481803500013336

Fernández-Delgado, M., Cernadas, E., Barro, S., & Amorim, D. (2014). Do we Need Hundreds of Classifiers to Solve Real World Classification Problems? The Journal of Machine Learning Research, 15(1), 3133–3181.

Fernandez Rojas, R., Hirachan, N., Brown, N., Waddington, G., Murtagh, L., Seymour, B., & Goecke, R. (2023). Multimodal physiological sensing for the assessment of acute pain. Frontiers in Pain Research, 4. 10.3389/fpain.2023.1150264

Fontana, M. A., Lyman, S., Sarker, G. K., Padgett, D. E., & MacLean, C. H. (2019). Can Machine Learning Algorithms Predict Which Patients Will Achieve Minimally Clinically Important Differences From Total Joint Arthroplasty? Clinical Orthopaedics & Related Research, 477(6), 1267–1279. 10.1097/CORR.0000000000000687

Furman, A. J., Prokhorenko, M., Keaser, M. L., Zhang, J., Chen, S., Mazaheri, A., & Seminowicz, D. A. (2020). Sensorimotor Peak Alpha Frequency Is a Reliable Biomarker of Prolonged Pain Sensitivity. Cerebral Cortex, 30(12), 6069–6082. 10.1093/cercor/bhaa124

Gabard-Durnam, L. J., Mendez Leal, A. S., Wilkinson, C. L., & Levin, A. R. (2018). The Harvard Automated Processing Pipeline for Electroencephalography (HAPPE): Standardized Processing Software for Developmental and High-Artifact Data. Frontiers in Neuroscience, 12. 10.3389/fnins.2018.00097

Garcia-Larrea, L., & Peyron, R. (2013). Pain matrices and neuropathic pain matrices: A review. Pain, 154, S29–S43. 10.1016/j.pain.2013.09.001

Gatchel, R. J., Peng, Y. B., Peters, M. L., Fuchs, P. N., & Turk, D. C. (2007). The biopsychosocial approach to chronic pain: Scientific advances and future directions. Psychological Bulletin, 133(4), 581–624. 10.1037/0033-2909.133.4.581

Géron, A. (2019). Hands-on Machine Learning with Scikit-Learn, Keras, and TensorFlow: Concepts, Tools, and Techniques to Build Intelligent (2nd ed.). O’Reilly.

Gers, F. A., Schmidhuber, J., & Cummins, F. (2000). Learning to Forget: Continual Prediction with LSTM. Neural Computation, 12(10), 2451–2471. 10.1162/089976600300015015

Gross, J., Schnitzler, A., Timmermann, L., & Ploner, M. (2007). Gamma Oscillations in Human Primary Somatosensory Cortex Reflect Pain Perception. PLoS Biology, 5(5), e133. 10.1371/journal.pbio.0050133

Grosse-Wentrup, M., Liefhold, C., Gramann, K., & Buss, M. (2009). Beamforming in Noninvasive Brain–Computer Interfaces. IEEE Transactions on Biomedical Engineering, 56(4), 1209–1219. 10.1109/TBME.2008.2009768

Herr, K., Coyne, P. J., McCaffery, M., Manworren, R., & Merkel, S. (2011). Pain Assessment in the Patient Unable to Self-Report: Position Statement with Clinical Practice Recommendations. Pain Management Nursing, 12(4), 230–250. 10.1016/j.pmn.2011.10.002

Hochreiter, S., & Schmidhuber, J. (1997). Long Short-Term Memory. Neural Computation, 9(8), 1735–1780. 10.1162/neco.1997.9.8.1735

Hsiao, F.-J., Chen, W.-T., Pan, L.-L. H., Liu, H.-Y., Wang, Y.-F., Chen, S.-P., Lai, K.-L., Coppola, G., & Wang, S.-J. (2022). Resting-state magnetoencephalographic oscillatory connectivity to identify patients with chronic migraine using machine learning. The Journal of Headache and Pain, 23(1), 130. 10.1186/s10194-022-01500-1

Huang, G., Xiao, P., Hung, Y. S., Iannetti, G. D., Zhang, Z. G., & Hu, L. (2013). A novel approach to predict subjective pain perception from single-trial laser-evoked potentials. NeuroImage, 81, 283–293. 10.1016/J.NEUROIMAGE.2013.05.017

Huang, Y., Li, W., Macheret, F., Gabriel, R. A., & Ohno-Machado, L. (2020). A tutorial on calibration measurements and calibration models for clinical prediction models. Journal of the American Medical Informatics Association, 27(4), 621–633. 10.1093/jamia/ocz228

Huber, M. T., Bartling, J., Pachur, D., Woikowsky-Biedau, S. v., & Lautenbacher, S. (2006). EEG responses to tonic heat pain. Experimental Brain Research, 173(1), 14–24. 10.1007/s00221-006-0366-1

Jensen, K. B., Regenbogen, C., Ohse, M. C., Frasnelli, J., Freiherr, J., & Lundström, J. N. (2016). Brain activations during pain. Pain, 157(6), 1279–1286. 10.1097/j.pain.0000000000000517

Jiang, T., Gradus, J. L., & Rosellini, A. J. (2020). Supervised Machine Learning: A Brief Primer. Behavior Therapy, 51(5), 675–687. 10.1016/j.beth.2020.05.002

Johnson, H. R., Trinidad, D. D., Guzman, S., Khan, Z., Parziale, J. V., DeBruyn, J. M., & Lents, N. H. (2016). A Machine Learning Approach for Using the Postmortem Skin Microbiome to Estimate the Postmortem Interval. PLOS ONE, 11(12), e0167370. 10.1371/journal.pone.0167370

Kaplan, A. Y., Fingelkurts, A. A., Fingelkurts, A. A., Borisov, S. V., & Darkhovsky, B. S. (2005). Nonstationary nature of the brain activity as revealed by EEG/MEG: Methodological, practical and conceptual challenges. Signal Processing, 85(11), 2190–2212. 10.1016/j.sigpro.2005.07.010

Keil, A., Bernat, E. M., Cohen, M. X., Ding, M., Fabiani, M., Gratton, G., Kappenman, E. S., Maris, E., Mathewson, K. E., Ward, R. T., & Weisz, N. (2022). Recommendations and publication guidelines for studies using frequency domain and time-frequency domain analyses of neural time series. Psychophysiology, 59(5). 10.1111/psyp.14052

Kim, J. A., & Davis, K. D. (2021). Neural Oscillations: Understanding a Neural Code of Pain. The Neuroscientist, 27(5), 544–570. 10.1177/1073858420958629

Kunz, M., Mylius, V., Scharmann, S., Schepelman, K., & Lautenbacher, S. (2009). Influence of dementia on multiple components of pain. European Journal of Pain, 13(3), 317–325. 10.1016/j.ejpain.2008.05.001

LeCun, Y., Bengio, Y., & Hinton, G. (2015). Deep learning. Nature, 521(7553), 436–444. 10.1038/nature14539

Ledowski, T., Bromilow, J., Paech, M. J., Storm, H., Hacking, R., & Schug, S. A. (2006). Monitoring of skin conductance to assess postoperative pain intensity. British Journal of Anaesthesia, 97(6), 862–865. 10.1093/bja/ael280

Lehmann, D. (1987). Principles of spatial analysis. In A. S. Gevins & A. Remond (Eds.), Handbook of electroencephalography and clinical neurophysiology: Methods of analysis of brain electrical and magnetic signals (pp. 309–354). Elsevier.

Lever, J., Krzywinski, M., & Altman, N. (2016). Model selection and overfitting. Nature Methods, 13(9), 703–704. 10.1038/nmeth.3968

Levi, D., Gispan, L., Giladi, N., & Fetaya, E. (2022). Evaluating and Calibrating Uncertainty Prediction in Regression Tasks. Sensors, 22(15), 5540. 10.3390/s22155540

Li, D., Puntillo, K., & Miaskowski, C. (2008). A Review of Objective Pain Measures for Use With Critical Care Adult Patients Unable to Self-Report. The Journal of Pain, 9(1), 2–10. 10.1016/j.jpain.2007.08.009

Li, L., Huang, G., Lin, Q., Liu, J., Zhang, S., & Zhang, Z. (2018). Magnitude and Temporal Variability of Inter-stimulus EEG Modulate the Linear Relationship Between Laser-Evoked Potentials and Fast-Pain Perception. Frontiers in Neuroscience, 12. 10.3389/fnins.2018.00340

Li, X., Zhang, S., Zhang, Q., Wei, X., Pan, Y., Zhao, J., Xin, X., Qin, C., Wang, X., Li, J., Yang, F., Zhao, Y., Yang, M., Wang, Q., Zheng, Z., Zheng, X., Yang, X., Whitlow, C. T., Gurcan, M. N., … Chen, K. (2019). Diagnosis of thyroid cancer using deep convolutional neural network models applied to sonographic images: a retrospective, multicohort, diagnostic study. The Lancet Oncology, 20(2), 193–201. 10.1016/S1470-2045(18)30762-9

Luo, W., Phung, D., Tran, T., Gupta, S., Rana, S., Karmakar, C., Shilton, A., Yearwood, J., Dimitrova, N., Ho, T. B., Venkatesh, S., & Berk, M. (2016). Guidelines for Developing and Reporting Machine Learning Predictive Models in Biomedical Research: A Multidisciplinary View. Journal of Medical Internet Research, 18(12), e323. 10.2196/jmir.5870

Mackey, S., Greely, H. T., & Martucci, K. T. (2019). Neuroimaging-based pain biomarkers: definitions, clinical and research applications, and evaluation frameworks to achieve personalized pain medicine. PAIN Reports, 4(4), e762. 10.1097/PR9.0000000000000762

Maniruzzaman, M., Rahman, M. J., Al-MehediHasan, M., Suri, H. S., Abedin, M. M., El-Baz, A., & Suri, J. S. (2018). Accurate Diabetes Risk Stratification Using Machine Learning: Role of Missing Value and Outliers. Journal of Medical Systems, 42(5), 92. 10.1007/s10916-018-0940-7

Marathe, A. R., Ries, A. J., & McDowell, K. (2014). Sliding HDCA: Single-Trial EEG Classification to Overcome and Quantify Temporal Variability. IEEE Transactions on Neural Systems and Rehabilitation Engineering, 22(2), 201–211. 10.1109/TNSRE.2014.2304884

Mari, T., Asgard, O., Henderson, J., Hewitt, D., Brown, C., Stancak, A., & Fallon, N. (2023). External validation of binary machine learning models for pain intensity perception classification from EEG in healthy individuals. Scientific Reports, 13(1), 242. 10.1038/s41598-022-27298-1

Mari, T., Henderson, J., Maden, M., Nevitt, S., Duarte, R., & Fallon, N. (2022). Systematic Review of the Effectiveness of Machine Learning Algorithms for Classifying Pain Intensity, Phenotype or Treatment Outcomes Using Electroencephalogram Data. The Journal of Pain, 23(3), 349–369. 10.1016/j.jpain.2021.07.011

McGuire, D. B., Kaiser, K. S., Haisfield-Wolfe, M. E., & Iyamu, F. (2016). Pain Assessment in Noncommunicative Adult Palliative Care Patients. Nursing Clinics of North America, 51(3), 397–431. 10.1016/j.cnur.2016.05.009

Miao, Y., Jin, J., Daly, I., Zuo, C., Wang, X., Cichocki, A., & Jung, T.-P. (2021). Learning Common Time-Frequency-Spatial Patterns for Motor Imagery Classification. IEEE Transactions on Neural Systems and Rehabilitation Engineering, 29, 699–707. 10.1109/TNSRE.2021.3071140

Michail, G., Dresel, C., Witkovský, V., Stankewitz, A., & Schulz, E. (2016). Neuronal oscillations in various frequency bands differ between pain and touch. Frontiers in Human Neuroscience, 10, 182.

Mienye, I. D., & Sun, Y. (2022). A Survey of Ensemble Learning: Concepts, Algorithms, Applications, and Prospects. IEEE Access, 10, 99129–99149. 10.1109/ACCESS.2022.3207287

Moon, S., Song, H.-J., Sharma, V. D., Lyons, K. E., Pahwa, R., Akinwuntan, A. E., & Devos, H. (2020). Classification of Parkinson’s disease and essential tremor based on balance and gait characteristics from wearable motion sensors via machine learning techniques: a data-driven approach. Journal of NeuroEngineering and Rehabilitation, 17(1), 125. 10.1186/s12984-020-00756-5

Moons, K. G. M., Altman, D. G., Reitsma, J. B., Ioannidis, J. P. A., Macaskill, P., Steyerberg, E. W., Vickers, A. J., Ransohoff, D. F., & Collins, G. S. (2015). Transparent Reporting of a multivariable prediction model for Individual Prognosis Or Diagnosis (TRIPOD): Explanation and Elaboration. Annals of Internal Medicine, 162(1), W1–W73. 10.7326/M14-0698

Mullen, T. (2012). CleanLine EEGLAB Plugin. San Diego, CA: Neuroimaging Informatic Tools and Resources Clearinghouse (NITRC).

Neuper, C., & Pfurtscheller, G. (2001). Event-related dynamics of cortical rhythms: frequency-specific features and functional correlates. International Journal of Psychophysiology, 43(1), 41–58. 10.1016/S0167-8760(01)00178-7

Okolo, C., & Omurtag, A. (2018). Research : Use of Dry Electroencephalogram and Support Vector for Objective Pain Assessment. Biomedical Instrumentation & Technology, 52(5), 372–378. 10.2345/0899-8205-52.5.372

Pedregosa, F., Varoquaux, G., Gramfort, A., Michel, V., Thirion, B., Grisel, O., Blondel, M., Prettenhofer, P., Weiss, R., Dubourg, V., Vanderplas, J., Passos, A., Cournapeau, D., Brucher, M., Perrot, M., & Duchesnay, E. (2011). Scikit-learn: Machine Learning in Python. Journal of Machine Learning Research, 12, 2825--2830.

Peirce, J. W. (2007). PsychoPy—Psychophysics software in Python. Journal of Neuroscience Methods, 162(1–2), 8–13. 10.1016/j.jneumeth.2006.11.017

Petre, B., Kragel, P., Atlas, L. Y., Geuter, S., Jepma, M., Koban, L., Krishnan, A., Lopez-Sola, M., Losin, E. A. R., Roy, M., Woo, C.-W., & Wager, T. D. (2022). A multistudy analysis reveals that evoked pain intensity representation is distributed across brain systems. PLOS Biology, 20(5), e3001620. 10.1371/journal.pbio.3001620

Peyron, R., Laurent, B., & García-Larrea, L. (2000). Functional imaging of brain responses to pain. A review and meta-analysis (2000). Neurophysiologie Clinique/Clinical Neurophysiology, 30(5), 263–288. 10.1016/S0987-7053(00)00227-6

Pfurtscheller, G. (1992). Event-related synchronization (ERS): an electrophysiological correlate of cortical areas at rest. Electroencephalography and Clinical Neurophysiology, 83(1), 62–69. 10.1016/0013-4694(92)90133-3

Pfurtscheller, G. (2001). Functional brain imaging based on ERD/ERS. Vision Research, 41(10–11), 1257–1260. 10.1016/S0042-6989(00)00235-2

Pfurtscheller, G., & Aranibar, A. (1977). Event-related cortical desynchronization detected by power measurements of scalp EEG. Electroencephalography and Clinical Neurophysiology, 42(6), 817–826. 10.1016/0013-4694(77)90235-8

Pfurtscheller, G., & Aranibar, A. (1979). Evaluation of event-related desynchronization (ERD) preceding and following voluntary self-paced movement. Electroencephalography and Clinical Neurophysiology, 46(2), 138–146. 10.1016/0013-4694(79)90063-4

Pfurtscheller, G., & Lopes da Silva, F. H. H. (1999). Event-related EEG/MEG synchronization and desynchronization: basic principles. Clinical Neurophysiology, 110(11), 1842–1857. 10.1016/S1388-2457(99)00141-8

Pfurtscheller, G., & Neuper, C. (1992). Simultaneous EEG 10 Hz desynchronization and 40 Hz synchronization during finger movements. NeuroReport, 3(12), 1057–1060. 10.1097/00001756-199212000-00006

Ploner, M., Sorg, C., & Gross, J. (2017). Brain Rhythms of Pain. Trends in Cognitive Sciences, 21(2), 100–110. 10.1016/j.tics.2016.12.001

Ramspek, C. L., Jager, K. J., Dekker, F. W., Zoccali, C., & van Diepen, M. (2021). External validation of prognostic models: what, why, how, when and where? Clinical Kidney Journal, 14(1), 49–58. 10.1093/ckj/sfaa188

Rivet, B., Souloumiac, A., Attina, V., & Gibert, G. (2009). xDAWN Algorithm to Enhance Evoked Potentials: Application to Brain–Computer Interface. IEEE Transactions on Biomedical Engineering, 56(8), 2035–2043. 10.1109/TBME.2009.2012869

Rockholt, M. M., Kenefati, G., Doan, L. V., Chen, Z. S., & Wang, J. (2023). In search of a composite biomarker for chronic pain by way of EEG and machine learning: where do we currently stand? Frontiers in Neuroscience, 17. 10.3389/fnins.2023.1186418

Sai, C. Y., Mokhtar, N., Yip, H. W., Bak, L. L. M., Hasan, M. S., Arof, H., Cumming, P., & Mat Adenan, N. A. (2019). Objective identification of pain due to uterine contraction during the first stage of labour using continuous EEG signals and SVM. Sādhanā, 44(4), 87. 10.1007/s12046-019-1058-4

Schiavenato, M., & Craig, K. D. (2010). Pain Assessment as a Social Transaction. The Clinical Journal of Pain, 26(8), 667–676. 10.1097/AJP.0b013e3181e72507

Schnakers, C., & Zasler, N. D. (2007). Pain assessment and management in disorders of consciousness. Current Opinion in Neurology, 20(6), 620–626. 10.1097/WCO.0b013e3282f169d9

Schomer, D. L., & Lopes, D. S. F. (2010). Niedermeyer’s electroencephalography : Basic principles, clinical applications, and related fields. Wolters Kluwer Health.

Schulz, E., Zherdin, A., Tiemann, L., Plant, C., & Ploner, M. (2012). Decoding an Individual’s Sensitivity to Pain from the Multivariate Analysis of EEG Data. Cerebral Cortex, 22, 1118–1123. 10.1093/cercor/bhr186

Shao, S., Shen, K., Yu, K., Wilder-Smith, E. P. V., & Li, X. (2012). Frequency-domain EEG source analysis for acute tonic cold pain perception. Clinical Neurophysiology, 123(10), 2042–2049. 10.1016/j.clinph.2012.02.084

Shirvalkar, P., Prosky, J., Chin, G., Ahmadipour, P., Sani, O. G., Desai, M., Schmitgen, A., Dawes, H., Shanechi, M. M., Starr, P. A., & Chang, E. F. (2023). First-in-human prediction of chronic pain state using intracranial neural biomarkers. Nature Neuroscience, 26(6), 1090–1099. 10.1038/s41593-023-01338-z

Singh, S. (2014). Magnetoencephalography: Basic principles. Annals of Indian Academy of Neurology, 17(5), 107. 10.4103/0972-2327.128676

Siontis, G. C. M., Tzoulaki, I., Castaldi, P. J., & Ioannidis, J. P. A. (2015). External validation of new risk prediction models is infrequent and reveals worse prognostic discrimination. Journal of Clinical Epidemiology, 68(1), 25–34. 10.1016/j.jclinepi.2014.09.007

Snell, K. I. E., Archer, L., Ensor, J., Bonnett, L. J., Debray, T. P. A., Phillips, B., Collins, G. S., & Riley, R. D. (2021). External validation of clinical prediction models: simulation-based sample size calculations were more reliable than rules-of-thumb. Journal of Clinical Epidemiology, 135, 79–89. 10.1016/j.jclinepi.2021.02.011

Sullivan, M. J. L., Bishop, S. R., & Pivik, J. (1995). The Pain Catastrophizing Scale: Development and validation. Psychological Assessment, 7(4), 524–532. 10.1037/1040-3590.7.4.524

Tivadar, R. I., & Murray, M. M. (2019). A Primer on Electroencephalography and Event-Related Potentials for Organizational Neuroscience. Organizational Research Methods, 22(1), 69–94. 10.1177/1094428118804657

Tracey, I., Woolf, C. J., & Andrews, N. A. (2019). Composite Pain Biomarker Signatures for Objective Assessment and Effective Treatment. Neuron, 101(5), 783–800. 10.1016/j.neuron.2019.02.019

Tu, Y., Tan, A., Bai, Y., Hung, Y. S., & Zhang, Z. (2016). Decoding Subjective Intensity of Nociceptive Pain from Pre-stimulus and Post-stimulus Brain Activities. Frontiers in Computational Neuroscience, 10. 10.3389/fncom.2016.00032

Vabalas, A., Gowen, E., Poliakoff, E., & Casson, A. J. (2019). Machine learning algorithm validation with a limited sample size. PLOS ONE, 14(11), e0224365. 10.1371/journal.pone.0224365

Van Calster, B., McLernon, D. J., van Smeden, M., Wynants, L., & Steyerberg, E. W. (2019). Calibration: the Achilles heel of predictive analytics. BMC Medicine, 17(1), 230. 10.1186/s12916-019-1466-7

Van Calster, B., Nieboer, D., Vergouwe, Y., De Cock, B., Pencina, M. J., & Steyerberg, E. W. (2016). A calibration hierarchy for risk models was defined: from utopia to empirical data. Journal of Clinical Epidemiology, 74, 167–176. 10.1016/j.jclinepi.2015.12.005

van der Miesen, M. M., Lindquist, M. A., & Wager, T. D. (2019). Neuroimaging-based biomarkers for pain. PAIN Reports, 4(4), e751. 10.1097/PR9.0000000000000751

Vargas-Lopez, O., Perez-Ramirez, C. A., Valtierra-Rodriguez, M., Yanez-Borjas, J. J., & Amezquita-Sanchez, J. P. (2021). An Explainable Machine Learning Approach Based on Statistical Indexes and SVM for Stress Detection in Automobile Drivers Using Electromyographic Signals. Sensors, 21(9), 3155. 10.3390/s21093155

Varma, S., & Simon, R. (2006). Bias in error estimation when using cross-validation for model selection. BMC Bioinformatics, 7(1), 91. 10.1186/1471-2105-7-91

Varoquaux, G. (2018). Cross-validation failure: Small sample sizes lead to large error bars. NeuroImage, 180, 68–77. 10.1016/j.neuroimage.2017.06.061

Vimala, V., Ramar, K., & Ettappan, M. (2019). An Intelligent Sleep Apnea Classification System Based on EEG Signals. Journal of Medical Systems, 43(2), 36. 10.1007/s10916-018-1146-8

Voepel-Lewis, T., Merkel, S., Tait, A. R., Trzcinka, A., & Malviya, S. (2002). The Reliability and Validity of the Face, Legs, Activity, Cry, Consolability Observational Tool as a Measure of Pain in Children with Cognitive Impairment. Anesthesia & Analgesia, 95(5), 1224– 1229. 10.1097/00000539-200211000-00020

Wager, T. D., Atlas, L. Y., Lindquist, M. A., Roy, M., Woo, C.-W., & Kross, E. (2013). An fMRI-Based Neurologic Signature of Physical Pain. New England Journal of Medicine, 368(15), 1388–1397. 10.1056/NEJMoa1204471

Watkinson, P., Wood, A. M., Lloyd, D. M., & Brown, G. D. A. (2013). Pain ratings reflect cognitive context: A range frequency model of pain perception. Pain, 154(5), 743–749. 10.1016/j.pain.2013.01.016

Willmott, C., & Matsuura, K. (2005). Advantages of the mean absolute error (MAE) over the root mean square error (RMSE) in assessing average model performance. Climate Research, 30, 79–82. 10.3354/cr030079

Witt, N., Coynor, S., Edwards, C., & Bradshaw, H. (2016). A Guide to Pain Assessment and Management in the Neonate. Current Emergency and Hospital Medicine Reports, 4(1), 1–10. 10.1007/s40138-016-0089-y

Xu, A., Larsen, B., Baller, E. B., Scott, J. C., Sharma, V., Adebimpe, A., Basbaum, A. I., Dworkin, R. H., Edwards, R. R., Woolf, C. J., Eickhoff, S. B., Eickhoff, C. R., & Satterthwaite, T. D. (2020). Convergent neural representations of experimentally-induced acute pain in healthy volunteers: A large-scale fMRI meta-analysis. Neuroscience & Biobehavioral Reviews, 112, 300–323. 10.1016/j.neubiorev.2020.01.004

Yang, L., & Shami, A. (2020). On hyperparameter optimization of machine learning algorithms: Theory and practice. Neurocomputing, 415, 295–316. 10.1016/j.neucom.2020.07.061

Yoshinaga, H., Nakahori, T., Ohtsuka, Y., Oka, E., Kitamura, Y., Kiriyama, H., Kinugasa, K., Miyamoto, K., & Hoshida, T. (2002). Benefit of Simultaneous Recording of EEG and MEG in Dipole Localization. Epilepsia, 43(8), 924–928. 10.1046/j.1528-1157.2002.42901.x

Younger, J., McCue, R., & Mackey, S. (2009). Pain outcomes: A brief review of instruments and techniques. Current Pain and Headache Reports, 13(1), 39–43. 10.1007/s11916-009-0009-x

Zhang, Z. G., Hu, L., Hung, Y. S., Mouraux, A., & Iannetti, G. D. (2012). Gamma-Band Oscillations in the Primary Somatosensory Cortex--A Direct and Obligatory Correlate of Subjective Pain Intensity. Journal of Neuroscience, 32(22), 7429–7438. 10.1523/JNEUROSCI.5877-11.2012

Zis, P., Liampas, A., Artemiadis, A., Tsalamandris, G., Neophytou, P., Unwin, Z., Kimiskidis, V. K., Hadjigeorgiou, G. M., Varrassi, G., Zhao, Y., & Sarrigiannis, P. G. (2022). EEG Recordings as Biomarkers of Pain Perception: Where Do We Stand and Where to Go? Pain and Therapy, 11(2), 369–380. 10.1007/s40122-022-00372-2

